# Human Lactoferrin is a Novel PFAS Target: Implications for Neo-natal Immune Function and Protein Stability

**DOI:** 10.64898/2026.08.25.746799

**Authors:** Mallory E. Thomas, Zachary S. McLean, Scott M. Belcher

## Abstract

Per-and polyfluoroalkyl substances (PFAS) constitute a diverse class of persistent synthetic chemicals utilized across industrial, medical, and consumer sectors that are pervasive global pollutants. Exposure to PFAS is linked to adverse impacts on both innate and adaptive immune systems. Human lactoferrin (hLF) is a key antimicrobial component of the developing innate immune system present in colostrum and breast milk. We hypothesized that hLF is a potential PFAS binding protein related to PFAS immunotoxicity. The results of thermal stability experiments indicated that all 11 tested PFAS bind and destabilize the structure of hLF. Notably PFBA, PFOS, HFPO-DA, and 6:2 FTSA decreased apo-hLF melting temperatures from 64°C to ≤ 37°C, suggesting that PFAS exposures destabilize the native hLF protein under physiological conditions. Relative binding affinities (K_d_) ranged from 0.2-11 mM across tested PFAS. Molecular docking was used to confirm experimental binding affinities and identify molecular interactions involved with PFAS binding. Calculated Gibbs Free Energies of binding ranged from −4.4 to −8.8 kcal/mol. Together, these results demonstrate that PFAS bind hLF at affinities comparable to human serum albumin and other PFAS binding proteins, and that some PFAS can destabilize hLF protein structure at physiologically relevant temperatures and conditions.

## Introduction

Per- and polyfluoroalkyl substances (PFAS) are highly fluorinated synthetic chemicals used widely across many industrial sectors with PFAS monomers and polymers routinely present in medical, pharmaceutical, consumer, and chemical products. The unique chemical properties of PFAS, including resistance to chemical and thermal degradation, and amphipathicity, have contributed to widespread global use. However, their degradation resistance, hydrophilic and oleophilic properties, have also contributed to persistent global PFAS contamination of the environment and biota ^1–3^. Previous evidence also links increased PFAS exposures with increased incidence of acute and long-term adverse health effects including cancers, liver and immune system dysfunction ^4–7^. Unlike lipophilic persistent organic pollutants, PFAS are proteophilic and primarily accumulate in well-perfuse protein-rich body compartments including blood, liver, and kidneys ^8–11^. While PFAS binding to serum and cellular transport proteins, albumin and liver fatty acid binding protein, is well-established, the diversity of other PFAS binding proteins, mechanisms of binding, and the functional consequences of PFAS/protein interactions remain poorly characterized ^12–14^.

Here we report findings of in vitro and in silico studies investigating PFAS binding to human lactoferrin (hLF), a multifunctional protein present at high concentrations in colostrum and breast milk ^15,16^. This research is an initial step in identifying potential PFAS binding proteins that have impacts on establishment protective neonatal immune functions. Because of the beneficial maternal contribution of hLF as a key immune-protective component of the immature neonatal immune system, we hypothesized that PFAS binding to hLF may serve as a key factor in maternal transfer and distribution of PFAS to neonates, and that PFAS binding could disrupt the antimicrobial immune activity of hLF ^17–19^. Furthermore, based on the substantial benefits of hLF on neonatal development, in particular hLF’s beneficial innate immune and anti-oxidant activities, hLF is also used as a dietary supplement in foods and infant formulas; PFAS contaminated hLF in these supplements might also be a previously unrecognized source of dietary PFAS exposure ^20^.

Human LF is an 80 kDa iron binding glycoprotein, composed of two conserved lobes of similar tertiary structures. Each lobe contains an identical ferric (Fe^3+^) binding site with iron coordinated by four amino acids: two tyrosines (Tyr92, Tyr192), asparagine (Asp60), and histidine (His253) in the N-lobe, and Tyr435, Tyr528, Asp395, and His597 in the C-lobe ^21–24^. The N-lobe (residues 1-333) and C-lobe (residues 345-694) are connected by a three-turn α-helix ^25,26^. Both Fe^+3^ binding sites also bind the counter ion carbonate (CO_3_^2-^) that binds prior to Fe^3+^ binding. Iron binding is cooperative with the C-lobe binding Fe^3+^ first to facilitate binding in N-lobe. Once binding occurs in both sites, a stable di-ferric “closed” conformation is formed ^27^. Because both lobes of hLF bind iron, hLF has three iron saturation forms that impact protein stability as indicated by 3 different melting temperatures (Tm) when analyzed by differential scanning calorimetry or fluorimetry (DSF). This characteristic Tm pattern corresponds to the high stability closed conformation (holo-hLF; Tm = 90°C); a minor single Fe^3+^ C-lobe bound conformation (Tm = 78°C), and Fe-free apo-conformation (Tm = 64°C) ^21^. In vivo, the physiological degree of iron saturation of native hLF is between 15-20% ^26,28–31,21^.

The aim of this study was to evaluate PFAS binding and define the relative binding affinity of hLF for a test set of 11 chemically and toxicologically informative PFAS using differential scanning fluorimetry (DSF), evaluate the impacts of PFAS binding on hLF protein stability, and computationally define molecular interactions governing PFAS binding to hLF and other transferrins.

## Materials and Methods

### Chemicals and Reagents

Human recombinant lactoferrin (90%, CAS 146897-68-9), ethylenediaminetetraacetic acid disodium salt dihydrate (EDTA, 99%, CAS 60-00-4), and 3-hydroxy-1,3-dimethyl-4(1H)- pyridine (DF, 98% purity, CAS 30652-11-0) were purchased from Millipore Sigma (Burlington, MA). GloMelt dye (λEx= 468 nm, λEm= 507 nm) was purchased from Biotium (Freemont, CA; Cat. No. 33022-1). Methanol (MeOH, 99.9% purity, Cat No. A454-4), dimethyl sulfoxide (DMSO, 99.9% purity, Cat. No. 136-1, Lot 147002), sodium phosphate dibasic (Na_2_HPO_4_) (99.2% purity, Cat. No. S374-3, Lot 056560), sodium chloride (100% purity, Cat. No. S271-10, Lot 134874), sodium bicarbonate (99% purity, CAS 144-55-8), and 4-(2-hydroxyethyl)-1-piperazineethanesulfonic acid (HEPES, 99% purity, Cat. No. BP310-1, Lot 052975) were purchased from ThermoFisher Scientific (Waltham, MA). Perfluorobutanoic acid (PFBA, 99% purity, CAS 375-22-4), perfluorooctanoic acid (PFOA, 95% purity, CAS 335-67-1), perfluoro(2-methyl-3-oxahexanoic) acid (HFPO-DA, 97% purity, CAS 13252-13-6) were purchased from Alfa Aesar (Havermill, MA). Perfluorohexanoic acid (PFHxA, 98% purity, CAS 307-24-4), perfluorobutanesulfonoic acid (PFBS, 98% purity, CAS 375-73-5) were purchased from TCI America (Portland, OR). Perfluorohexanesulfonic acid, (PFHxS, 98% purity, CAS 3871-99-6) was purchased from Frontier Scientific (Newark, DE). Perfluorooctanesulfonate, (PFOS, 98% purity, CAS 2795-39-3) and perfluoro-3,6,9-trioxadecanoic acid (PFO3DoDa, 98% purity, CAS 151772-59-7) were purchased from Matrix Scientific (Columbia, SC). Trifluoroacetic acid, (TFA, 99% purity, CAS 76-05-1) was purchased from Fisher Scientific (Waltham, MA). Lastly, 2H, 2H, 3H, 3H-perfluorooctane sulfonate, (6:2 FTSA, 98% purity, CAS 59587-38-1) and nonafluoro-3,6-dioxaheptanoic acid (PFDHA, 99% purity, CAS 151772-58-6) were from Synquest Laboratories (Alachua, FL).

### Chemical Preparation

Aqueous solutions were prepared with Milli-Q A10 water (18 Ω; 3 ppb total oxidizable organics). Stock solutions of hLF (0.20 mM), ethylenediaminetetraacetic acid (EDTA, 100 mM), deferiprone (DF, 250 mM), sodium bicarbonate (50 mM), and each PFAS (5, 10, or 20 mM) were prepared in aqueous HEPES-buffered saline (HBS; 140 mM sodium chloride, 50 mM HEPES, and 0.38 mM sodium phosphate dibasic, pH 7.38). Stocks of 50% DMSO and 50% MeOH were also made in 1X HBS. Desired working stock solutions were made by serial dilution.

### Differential Scanning Fluorimetry

All DSF experiments followed detailed published protocols ^32,33^. Assays were ran using a sealed optical 96-well reaction plate (MicroAmp Fast, Applied Biosystems) with a final volume of 20 µL per well ^32,34^. A minimum of two plates were run for each hLF-PFAS combination, with 4 replicates per concentration on each plate. Controls included: no protein negative control (NPC), vehicle controls, and three concentrations of PFHxA (0.3, 3, and 6 mM) as positive controls with all controls ran in quadruplicate. All hLF containing experimental and control samples had a final protein concentration of 0.02 mM in each well to yield an optimized signal to noise ratio ^34^.

### Molecular Docking

Autodock Vina (v.1.2.0) was employed for molecular docking analyses ^35,36^. The RCSB Protein Data Bank (PDB, https://www.rcsb.org) was used to select high resolution (≤ 2.5 Å) three-dimensional protein crystal structures. All PFAS ligand files were downloaded as three-dimensional SDF conformer files from the NCBI PubChem database (https://pubchem.ncbi.nlm.nih.gov/). Those files were converted to .mol2 files using PyMol (v.3.1)^37^ and prepared for docking using AutoDock Tools (v.1.5.7) ^36,38^. Molecular docking was utilized to identify preferential binding locations of PFAS to PDB crystal structure 1CB6 (apo-hLF) and holo-hLF (1LFG) for human lactoferrin ^22,23^. Carbonate ions remained in the structure, and the Fe^3+^ cations were removed from the holo-hLF structure. The Grid Box feature in AutoDock tools was used to determine the grid box dimensions for all analyzed structures (Table S1). Five replicates of each molecular docking simulation were run for all combinations utilizing an exhaustiveness level of 128 to ensure reliability of the search algorithm to find the highest affinity conformation.

Utilizing the holo-hLF crystal structure, dual ligand molecular docking was performed using AutoDockTools (v.1.2.7) using two ligand structures of both PFBA and PFBS. The grid box was localized around the carbonate pocket of the C-lobe with the following dimensions: x center = 38.01, y center = 28.137, z center = −17.573, and X= 12 Å, Y= 18 Å, and Z= 12 Å. Replicates and exhaustiveness parameters were identical to single ligand docking.

### Docking to other Transferrins

Protein crystal structures 1IQ7 (ovotransferrin) from *Gallus gallus*, 6XR0 (human melanotransferrin), 1NKX (bovine lactoferrin), and 4H0W (human serum transferrin) were employed for molecular docking to evaluate PFAS binding across the transferrin family and various species ^39–42^. In each case, the C-terminal lobe of the protein was analyzed for PFAS binding. Human melanotransferrin crystal structure 6XR0 required removal of complexed SC57.32 Fab region prior to docking experiments. Grid box dimensions, species information, and crystal structure PDB codes for each transferrin are provided in Table S1.

### Data Analysis

All DSF analysis was previously reported in detailed step-by-step protocols with the exception of data being collected beginning at 22^0^C ^33,34^. Melting temperature (Tm) values were determined using Protein Thermal Shift Software (v2.3, Applied Biosystems) with manual regions of analysis employed ^43^. All statistical and concentration-response analysis was done using Prism (v.10, GraphPad Software Inc.). The Tm was defined as the temperature at which the largest change in fluorescence per unit time was observed ^34^. Curve smoothing for concentration-response curves used the Savitzky and Golay method ^44^. Simple cooperative ligand binding models were used to calculate dissociation constants (K_d_) for all hLF-ligand combinations.

Utilizing standard term weights, the scoring function of AutoDock Vina was used to calculate ΔG_bind_ predictions of the top nine conformations of ligand binding for all hLF-PFAS combinations. Additionally, molecular docking output was analyzed for the possibility of hydrogen bond interactions using PyMOL’s putative polar contact function. The maximum hydrogen bond distance was 3.6 Å with a maximum default angle of 63° for assessing putative contacts.

## Results

### DSF validation

Using DSF, we confirmed the presence of three Tm values (Tm =64.5° ± 0.07, 77.2 ° ± 0.08°, and 89.1° ± 0.09° C; n = 15) corresponding to each of the three Fe^3+^ saturated states of hLF (Fig. 1A). To further confirm these peak assignments, concentration response analysis was conducted using the iron chelators deferiprone (DF) (Fig. 1B) or EDTA (Fig. S2). Increasing concentrations of both DF or EDTA resulted in elimination of peaks 2 and 3, with a notable decrease in the Tm for peak 1 (ΔTm = −4° C), which is considered due to a destabilizing impact from sequestration of all Fe^3+^ in equilibrium with hLF, and/or non-selective ion sequestration resulting from high concentrations of DF (Fig. 1B).

**Figure 1:**
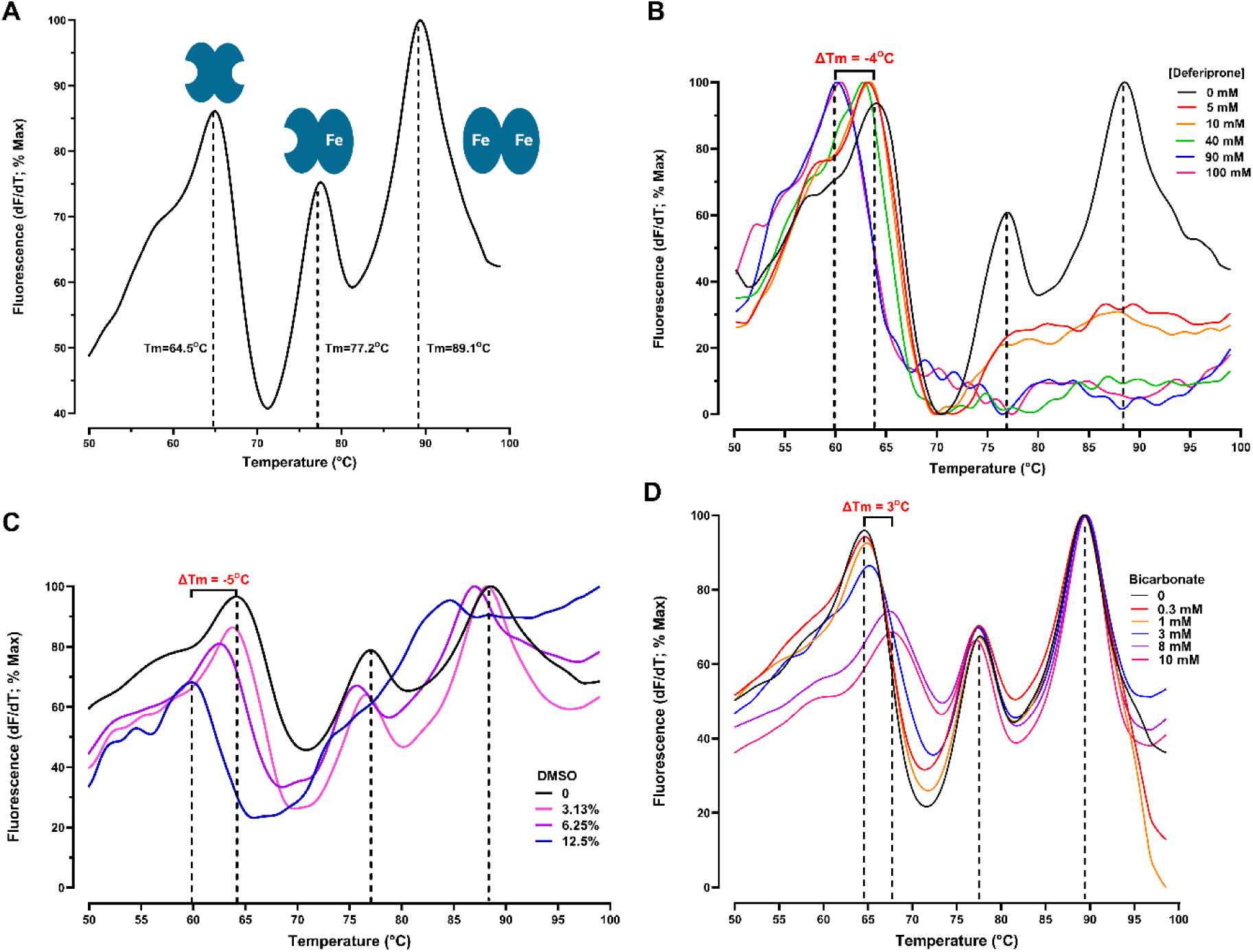
Control hLF DSF Experiments. (A) Normalized first derivative (df/dT) DSF spectrum for hLF. The Tm is denoted by dotted lines. Each conformation of hLF is denoted with each peak. (B) Normalized first derivative DSF spectrum for hLF with increasing additions of deferiprone. The change in Tm is defined, and the fluorescence maxima are denoted by dotted lines. (C) Normalized first derivative DSF spectrum for hLF with increasing percentages (v/v%) of DMSO with fluorescence maxima denoted by dotted lines. (D) Normalized first derivative DSF spectrum for hLF with increasing concentrations of sodium bicarbonate, with fluorescence maxima of hLF denoted by dotted lines.

Concentration response analysis was also performed to evaluate PFAS co-solvent effects of DMSO (Fig. 1C) or MeOH (Fig. S3). Each cosolvent decreased stability of apo-hLF at concentrations higher than 6.25% (*v/v*) with complete denaturation at 25% (*v/v*; not shown). The EC_50_ for the effect of DMSO and MeOH was 15.4% (*v/v*; r^2^ = 0.87) and 12.9% (*v/v;* r^2^ = 0.99), respectively. Therefore, we excluded cosolvents from all further experiments which limited the ability to evaluate long-chain congeners > C8, and fluorotelomer alcohols: only acidic forms of PFAS (PFCA, PFSA, perfluorinated alkyl ethers, and fluorotelomer sulfonic acids) were utilized in the study. Increasing concentrations of HCO_3_^-2^ resulted in a modest stabilizing effect on apo-hLF structure (Fig. 1D; ΔTm = 3° C ± 1, EC_50_ 5.5 mM, r^2^ = 0.94).

### Lactoferrin Binding Affinity for PFAS

Binding of each PFAS caused a destabilization of hLF as indicated by large decreases in Tm (Tm range = 30°C - 61°C) with remarkably large ΔTm values ranging from −2.8°C to −42°C (Fig. 2). Calculated EC_50_ and K_d_ values ranged from 0.2 - 11.0 mM (Table 1). Clear cooperative binding was observed with Hill Slopes >1 (Table 1) and cooperativity of PFAS binding was independent of bicarbonate.

**Figure 2:**
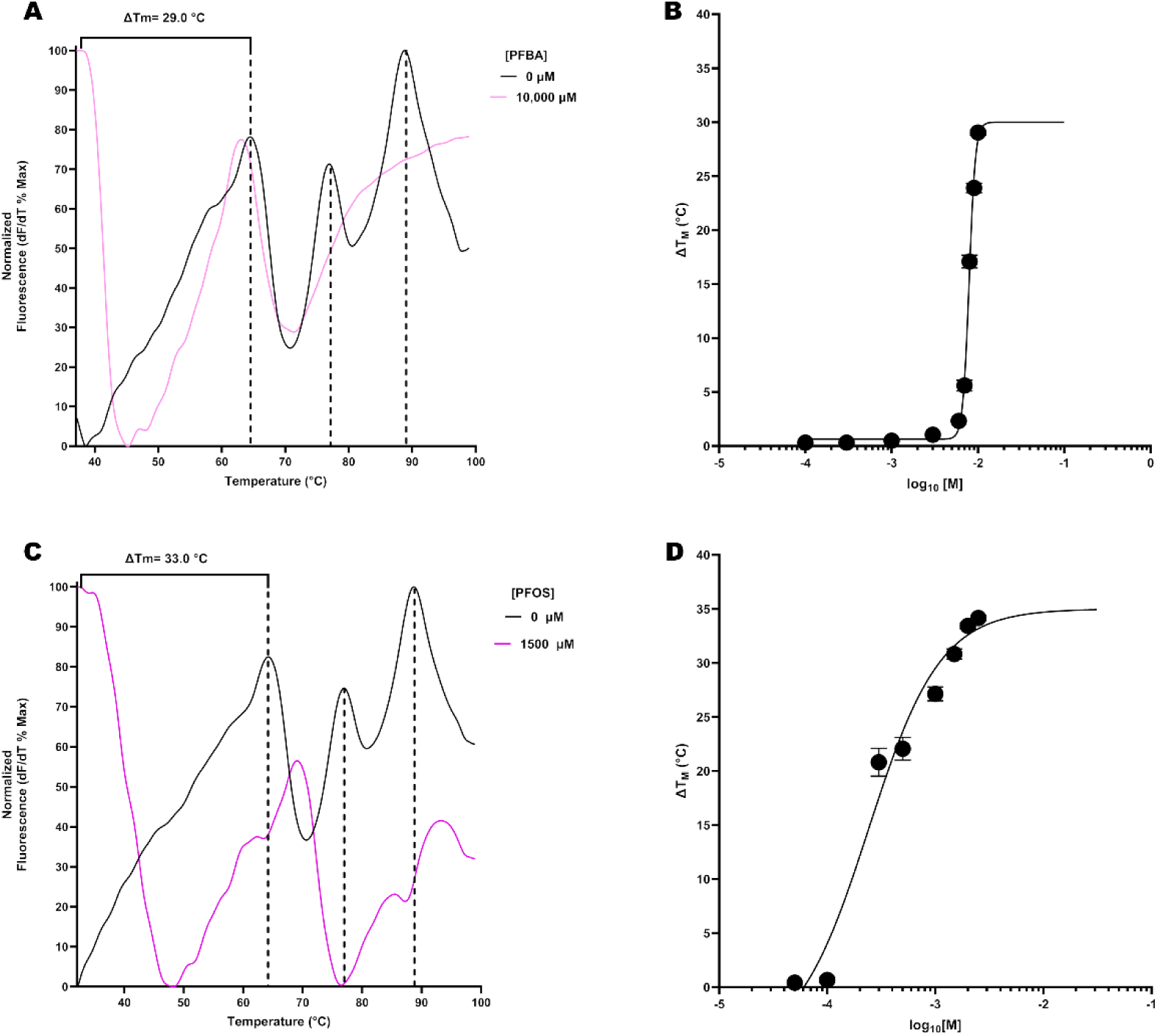
hLF DSF Experiments with PFBA and PFOS. (A) Normalized derivative fluorescence (df/dT) of hLF with increasing additions of PFBA. The fluorescence maxima values are denoted by dotted lines, with ΔTm depicted with a bracket. (B) Concentration response curve of hLF with increasing additions of PFBA where error bars indicate the standard error of the mean. (C) Normalized fluorescence of hLF with increasing additions of PFOS. The fluorescence maxima of hLF are denoted by dotted lines with ΔTm depicted with a bracket. (D) Concentration response curve of hLF with increasing additions of PFOS. The error bars indicate the standard error of the mean.

**Table 1:** hLF Binding Affinities for PFAS.

|  |  |  | No Bicarbonate |  |  |  | 8.0 mM Bicarbonate |  |  |  |
| --- | --- | --- | --- | --- | --- | --- | --- | --- | --- | --- |
| PFAS | C<sub>n</sub>F<sub>2/3</sub> | F<sub>n</sub> | R<sup>2</sup> | Max ΔT<sub>M</sub> | K<sub>d</sub> | HS | R<sup>2</sup> | Max ΔT<sub>M</sub> | K<sub>d</sub> | HS |
| Perfluorinated Carboxylic Acids |  |  |  |  |  |  |  |  |  |  |
| TFA | 2 | 3 | 0.96 | -3.8±0.1 | 7.7±0.1 | 9.3 | 0.86 | -4.0±0.6 | 15.0±0.3 | 4.1 |
| PFBA | 4 | 7 | 0.99 | -29.0±0.2 | 7.9±0.0 | 11.9 | 0.82 | -1.8±0.2 | 8.7±0.5 | 1.6 |
| PFHxA | 6 | 11 | 0.83 | -2.7±0.2 | 4.7±0.3 | 2.7 | 0.94 | -2.9±0.1 | 4.0±0.1 | 2.3 |
| PFOA | 8 | 15 | 0.99 | -42.8±0.1 | 0.6±0.01 | 5.3 | 0.99 | -42.7±0.6 | 0.6±0.01 | 9.4 |
| Perfluorinated Sulfonic Acids |  |  |  |  |  |  |  |  |  |  |
| PFBS | 4 | 9 | 0.97 | -8.4±0.4 | 7.7±0.4 | 3.0 | 0.94 | -6.0±0.6 | 7.0±0.2 | 3.2 |
| PFHxS | 6 | 13 | 0.91 | -3.2±0.2 | 0.6±0.0 | 5.9 | - | - | - | - |
| PFOS | 8 | 17 | 0.96 | -34.2±0.3 | 0.2±0.0 | 1.3 | 0.98 | -32.9±0.5 | 0.4±0.0 | 2.7 |
| Perfluorinated Alkyl Ethers |  |  |  |  |  |  |  |  |  |  |
| PFDHA | 5 | 9 | 0.94 | -18.5±2.5 | 11.0±0.1 | 12.1 | - | - | - | - |
| HFPO-DA | 5 | 11 | 0.95 | -28.6±0.8 | 4.6±0.2 | 1.9 | - | - | - | - |
| PFO3DoDa | 7 | 13 | 0.76 | -3.1±0.5 | 0.4±0.0 | 6.8 | - | - | - | - |
| Fluorotelomer Sulfonic Acid |  |  |  |  |  |  |  |  |  |  |
| 6:2 FTSA | 6 | 13 | 0.97 | -31.4±0.2 | 0.3±0.0 | 4.6 | - | - | - | - |

C<sub>n</sub>F<sub>2/3</sub> is the number of aliphatic carbons, F<sub>n</sub> is the number of fluorines, $\Delta T_M$ is in °C, K<sub>d</sub> is reported as mean values with ± SEM in mM, and HS is the Hill Slope. Each compound was analyzed on at least two separate plates with n ≥ 4 per plate. Hyphens indicate compounds where analysis was not performed.

Perfluoroalkyl sulfonic acids were bound with higher affinity than perfluoroalkyl carboxylic acids and increasing alkyl chain length was associated with an increase in binding affinity (Table 1). The presence of bicarbonate had minimal impacts on K_d_ values for each PFAS examined except for TFA where the binding affinity was decreased by 51% (Table 1).

### Molecular Docking analysis

“Blind docking” simulations for full length apo-hLF revealed that strongest PFAS binding was localized at the C-terminal Fe^3+^ binding site for all PFAS. Initial simulations of PFAS to the full length “open” apo-hLF and the “closed” holo-conformation identified only minor differences in calculated ΔG_bind_. Differences in ΔG_bind_ increased with increasing chain length for PFCAs with the ΔΔG ranging from 0.1 kcal/mol for TFA and PFBA to 0.9 kcal/mol for PFOA (Table 2). When docking was performed for the N-lobe and the C-lobes of the “open” apo conformation only very small differences were observed (*M* = 0.24 kcal/mol; max = 0.7 kcal/mol for PFHxA; n = 9). For the holo-conformation there was clear preferential binding at the C-lobe; a minimum difference in ΔG_bind_ of 1.0 kcal/mol for TFA and a maximal differencev3.0 kcal/mol for HFPO-DA and PFO3DoDa was observed. (Table 2).

**Table 2.** Delta G of binding estimates from molecular docking analysis.

| | $C_nF_{2/3}$ | $F_n$ | 1CB6 | C-Lobe | N-lobe | 1LFG | C-Lobe | N-lobe |
| --- | --- | --- | --- | --- | --- | --- | --- | --- |
| **PFCA** | | | $\Delta G_{\text{bind}}$ | | | $\Delta G_{\text{bind}}$ | | |
| TFA | 2 | 3 | -4.5 | -4.4 | -4.5 | -4.4 | -4.4 | -3.5 |
| PFBA | 4 | 7 | -6.4 | -6.0 | -6.0 | -6.5 | -6.6 | -4.5 |
| PFHxA | 6 | 11 | -7.7 | -7.7 | -7.0 | -8.2 | -8.0 | -5.6 |
| PFOA | 8 | 15 | -8.0 | -8.3 | -8.0 | -8.9 | -8.4 | -5.9 |
| **PFSA** |  |  |  |  |  |  |  |  |
| PFBS | 4 | 9 | -7.2 | -7.2 | -6.9 | -7.6 | -7.6 | -5.1 |
| PFHxS | 6 | 13 | -7.7 | -7.9 | -7.7 | -8.9 | -7.9 | -5.7 |
| PFOS | 8 | 17 | -8.7 | -8.6 | -8.6 | -9.8 | -8.7 | -6.1 |
| **PFEA** |  |  |  |  |  |  |  |  |
| HFPO-DA | 5 | 11 | -7.7 | -7.7 | -7.4 | -7.9 | -8.0 | -5.0 |
| PFDHA | 5 | 9 | -7.5 | -7.4 | -7.5 | -7.7 | -7.5 | -5.3 |
| PFO3DoDa | 7 | 13 | -8.3 | -8.3 | -8.2 | -8.8 | -8.6 | -5.6 |
| **PFTSA** |  |  |  |  |  |  |  |  |
| 6:2 FTSA | 6 | 13 | -7.9 | -8.1 | -7.7 | -8.6 | -8.4 | -5.7 |

$C_nF_{2/3}$ is the number of aliphatic carbons, $F_n$ is the number of fluorines, and $\Delta G_{\text{bind}}$ refers to the Gibbs Free energy of binding predicted by molecular docking for the most spontaneous conformation of the ligand for hLF binding. 1CB6 is the apo-hLF conformer, and 1LFG is the holo-hLF conformer.

For many tested PFAS, binding involved hydrogen bonding with amino acids Arg465, Asp644, and His597. Interactions to these amino acids were proximal to the C-lobe bicarbonate solvent Fe^3+^ binding pocket of holo-hLF. Residue Arg465, which interacts with bicarbonate in the iron binding pocket, was found to have direct hydrogen bond interactions with PFOS, PFHxS, HFPO-DA, PFO3DoDa, and 6:2 FTSA. Even without bicarbonate present, PFHxA, PFBA, PFBS, PFDHA, and PFOA interacted via hydrogen bonding to His597 (Fig. 3), and TFA was predicted to hydrogen bond with Arg587 near the bicarbonate binding pocket.

**Figure 3:**
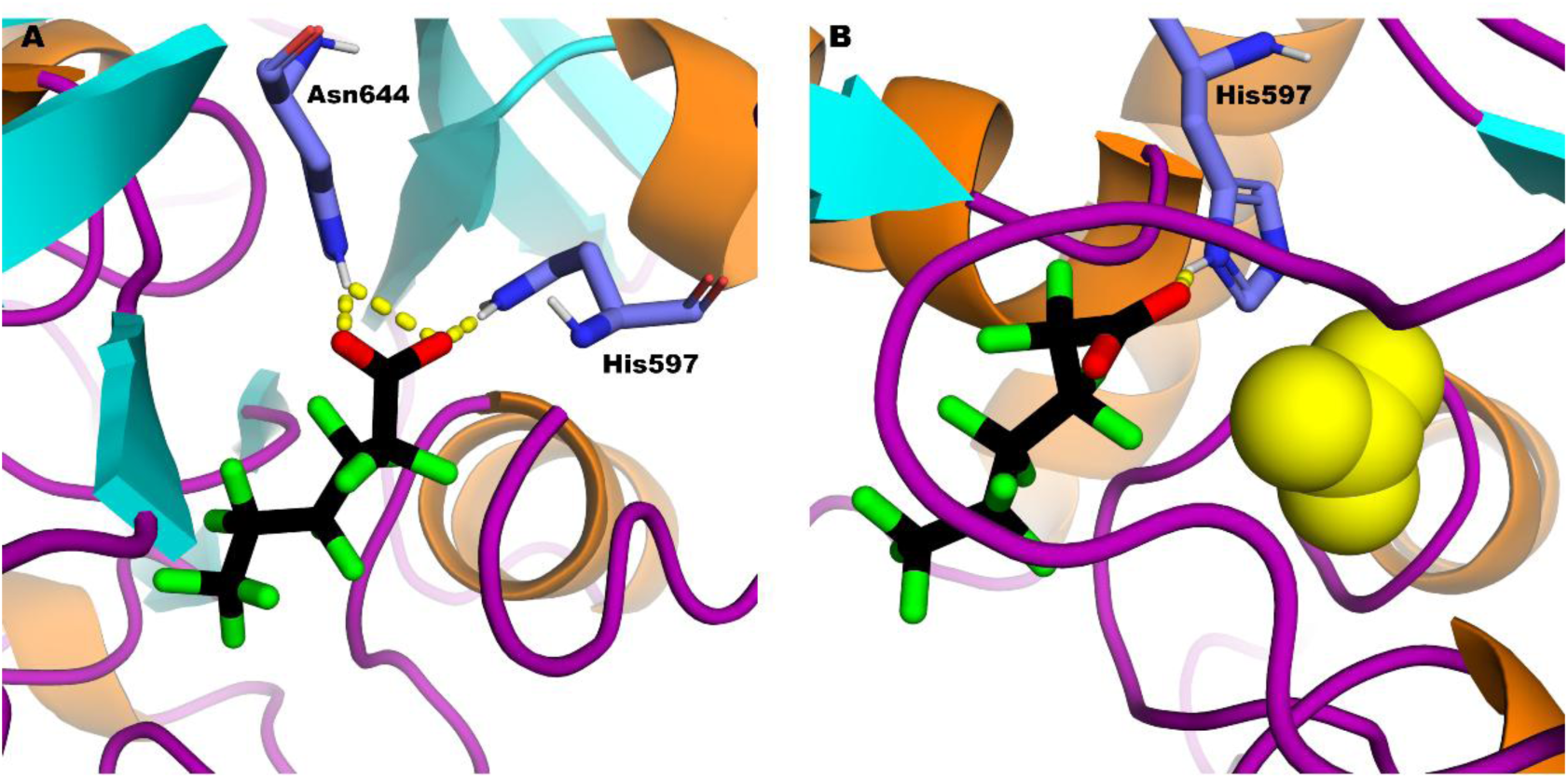
Molecular Docking Prediction of PFHxA with holo-hLF and apo-hLF. Ribbon diagrams of apo-hLF (A) and holo-hLF (B) with the most spontaneous conformation of PFHxA are shown with hydrogen bonds depicted as dashed lines. The amino acids involved in hydrogen bonding are shown in stick representations and labeled. In holo-hLF (B) the carbonate anion is depicted as space filling spheres.

Binding with the N-terminal lobe of apo-hLF was associated with a decrease in spontaneity of PFAS binding for PFHxA, PFOA, PFBS, PFHxS, HFPO-DA, PFO3DoDa, and 6:2 FTSA as indicated by their less negative ΔG_bind_ (Table 2). A majority of PFAS were predicted to form hydrogen bonds adjacent to the bicarbonate binding pocket in N-lobe residues Arg249 and His246. TFA was the only PFAS not predicted to form hydrogen bond interactions (Table 3).

**Table 3:** C-lobe molecular docking results to apo-hLF and holo-hLF.

| Ligand | 1CB6 |  | 1LFG |  |
| --- | --- | --- | --- | --- |
|  | # H Bonds | Residues | # H Bonds | Residues |
| **PFCA** |  |  |  |  |
| TFA | 1 | Arg587 | 2 | Asn553, Asn640 |
| PFBA | 3 | His597, Asn644 | 1 | His597 |
| PFHxA | 3 | His597, Asn644 | 1 | His597 |
| PFOA | 3 | His597, Asn644 | 0 | - |
| **PFSA** |  |  |  |  |
| PFBS | 3 | His597, Asn644 | 0 | - |
| PFHxS | 2 | Asp394, Arg465 | 2 | Arg533, Tyr562 |
| PFOS | 3 | Arg465 | 1 | Glu511 |
| **PFEA** |  |  |  |  |
| HFPO-DA | 5 | Arg465, Asn644 | 0 | - |
| PFDHA | 5 | Arg465, Tyr526, His597, Asn644 | 3 | Arg465 |
| PFO3DoDa | 4 | Arg465, Tyr526, Thr527, Asn644 | 2 | Arg465 |
| **PFTSA** |  |  |  |  |
| 6:2 FTSA | 4 | Asn368, Arg465, Tyr528 | 4 | Arg444, Asn640 |

Dashes indicate no hydrogen bonds.

### Docking to holo-hLF

Blind docking with full length holo-hLF structure revealed no preference for the C or N terminal binding sites. All PFAS examined were predicted to bind at the C-terminal Fe^3+^ binding pocket with lower affinity than without the carbonate ion present, see Table 2. The binding of PFHxA in both apo- and holo-conformers is shown in Figure 3.

Binding in the N-lobe of 1LFG revealed preference to locations adjacent to the carbonate binding pocket with five PFAS (PFBA, PFBS, 6:2 FTSA, PFDHA, and PFO3DoDa) illustrating hydrogen bond interactions with Arg224, close in proximity to carbonate binding amino acid of the N-lobe, His252 (Table 3). All 11 of the PFAS tested illustrated an increase in ΔG_bind_, and thus overall decreased affinity compared to the blind docking of holo-hLF with values ranging from −3.9 to −6.1 kcal/mol. PFHxA, PFHxS, PFOA, PFOS, and TFA illustrated hydrogen bond interactions to Lys101. HFPO-DA was the only PFAS to not exhibit hydrogen bond interactions, indicating binding is primarily mediated through van der Waals forces and hydrophobic interactions. In all cases, these amino acid residues lie outside of the iron-bicarbonate binding pocket.

All PFAS were bound in the C-terminal of holo-hLF with lower affinity than without the bicarbonate ion present, see Table 2. Molecular docking revealed a decrease in ΔG_bind_ of PFAS in the C-lobe of holo-hLF with energies ranging from −4.4 to −8.7 kcal/mol (Table 2). Docking of 4 PFAS, 6:2 FTSA, PFDHA, PFO3DoDa, and PFHxS, identified hydrogen bond interactions with arginine residues (Arg444, Arg465, and/or Arg553). The 3 strongest binding conformations for HFPO-DA, PFBS, and PFOA did not involve hydrogen bond interactions, and both PFBA and PFHxA revealed hydrogen bond interactions to His597, an amino acid responsible for bicarbonate-iron binding in the C-lobe (Table 3). Approximately half of the PFAS analyzed revealed either no change in Gibbs Free Energy, or a decrease in energy compared to that of the blind docking of holo-hLF. The remaining half showed slight increases in ΔG_bind_, and therefore a decreased affinity of binding (Table 2).

### PFAS Binding to Related Transferrins

Molecular docking of PFBA and PFHxA was performed for ovotransferrin from *Gallus gallus* (o-TF, PDB ID: 1IQ7), human melanotransferrin (m-TF, PDB ID: 6XR0), bovine lactoferrin (b-LF, PDB ID: 1NKX), and human serum transferrin (h-TF, PDB ID: 4H0W) protein structures (Table S1). RMSD values of protein alignment to apo-hLF revealed values of 0.778 for o-TF, 1.513 for m-TF, 4.520 for b-LF, and 7.167 for h-TF. Calculated ΔG_bind_ for PFBA and PFHxA are shown in Table 4. Docking simulations with PFBA revealed that binding was mediated mainly by van der Waals and hydrophobic interactions as no hydrogen bonding was identified. For h-TF binding of PFBA involved hydrogen bond interactions at The667 and Ser668, and for b-LF binding involved hydrogen bonding at Asp392 and His595. Binding of PFHxA was predicted to form hydrogen bond interactions with His625 of m-TF, and for o-TF hydrogen bonding was predicted at Arg648. In the case of h-TF and b-LF, hydrogen bond interactions were predicted to involve His585 and His595 respectively (Table 4. Fig. 4).

**Table 4:** Molecular docking to the transferrin family proteins.

| Protein | PDB ID | Ligand | #H Bonds | Residues | $\Delta G_{\text{bind}}$ |
| --- | --- | --- | --- | --- | --- |
| Ovotransferrin | 1IQ7 | PFBA | 0 | - | -5.7 |
|  |  | PFHxA | 2 | Arg648 | -6.4 |
| Melanotransferrin | 6XR0 | PFBA | 0 | - | -5.5 |
|  |  | PFHxA | 1 | His625 | -6.2 |
| Bovine Lactoferrin | 1NKX | PFBA | 2 | Asp392, His595 | -6.3 |
|  |  | PFHxA | 2 | Asp392, His595 | -7.9 |
| Serum Transferrin | 4H0W | PFBA | 2 | The667, Ser668 | -6.4 |
|  |  | PFHxA | 2 | His585 | -8.0 |

$\Delta G_{\text{bind}}$ is in kcal/mol. Dashes indicate no hydrogen bond interactions.

**Figure 4:**
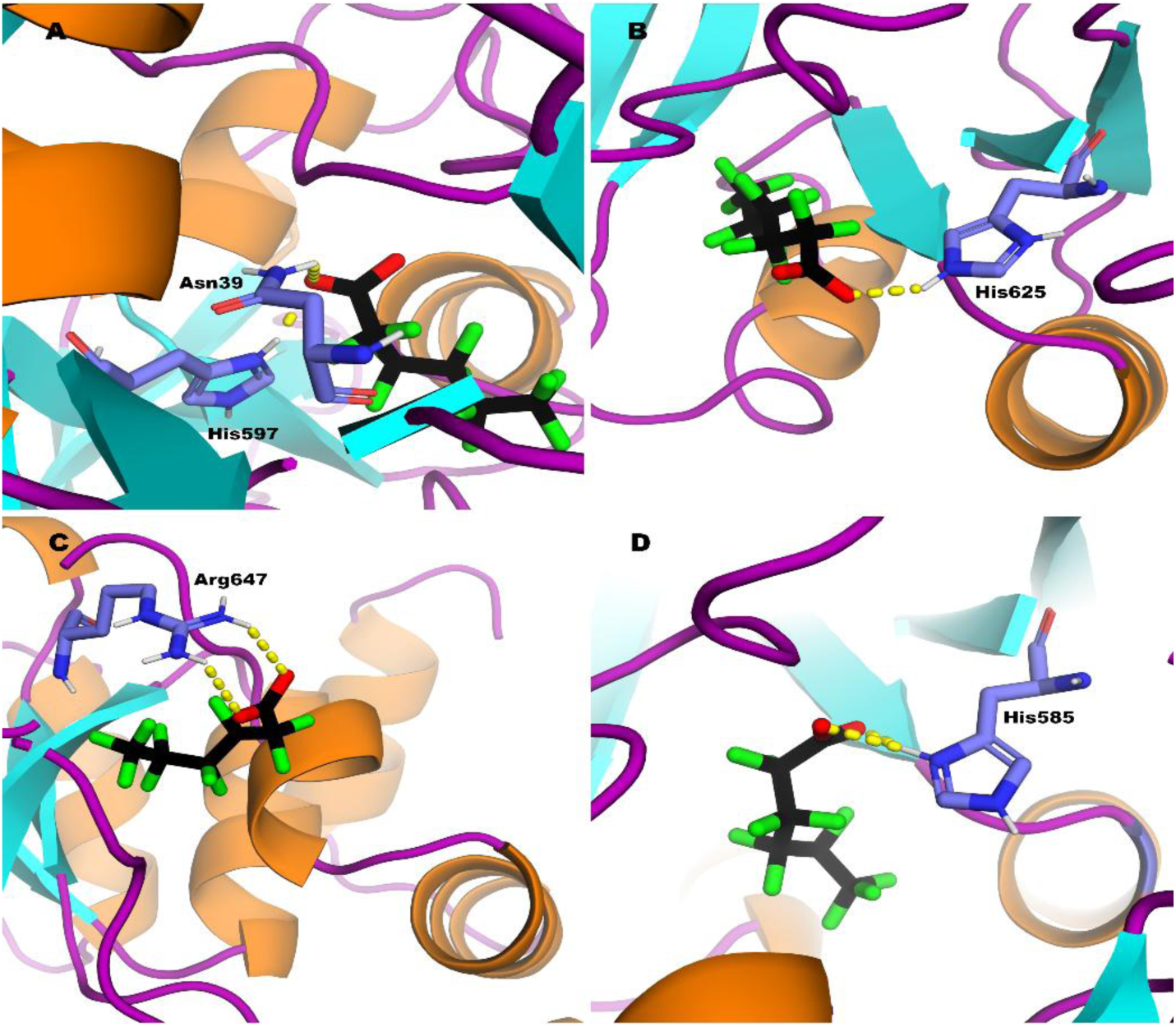
Molecular Docking Predictions of PFHxA to the Transferrin Family Proteins. Ribbon diagrams of b-LF (A), m-TF (B), o-TF (C), and h-TF (D) with the most spontaneous conformation of PFHxA are shown with hydrogen bonds depicted as dashed lines. The amino acids involved in hydrogen bonding are shown in stick representations and labeled.

## Discussion

Human breast milk and dairy products are well-established routes of PFAS exposure, particularly during infancy when the immune system is highly sensitive to toxicant exposures ^45–47^. Epidemiological studies have associated developmental PFAS exposure with immunosuppression, including diminished antibody responses to vaccinations ^48^. Here, we identified hLF and other transferrins as PFAS-binding proteins, suggesting that lactational exposure may not only transfer PFAS to neonates, but may also alter the function of these immunoprotective milk proteins. Because hLF is abundant in colostrum and mature breast milk, where it provides essential immune protection during neonatal immune development, PFAS induced destabilization of hLF protein structure could weaken the immunoprotective benefits of breast feeding ^15,16,49^. More broadly, PFAS binding across the transferrin family suggests a potential mechanism by which these chemicals can disrupt nutritional immunity, potentially contributing to the immunosuppression associated with PFAS exposure.

While PFAS binding to serum and cellular transport proteins, albumin and liver fatty acid binding protein is well-established, the diversity of other PFAS binding proteins, mechanisms of binding, and the functional consequences of PFAS/protein interactions remain poorly characterized ^12–14^. Molecular docking has been utilized as a robust method to suggest structural influences of PFAS and identify modes of binding across evolutionary conserved binding domains. Our results presented here-in have identified hLF and other transferrins as PFAS-binding proteins and suggest that adverse effects of PFAS-binding disorder native structure as indicated experimentally by decreased Tm. Regardless of carbon chain length, functional group, or ether oxygen content, all PFAS congeners tested significantly decreased hLF thermal stability. The calculated binding affinity and strength of binding was increased with increasing carbon chain length and was generally higher for PFSAs than PFCAs. Short chain and ultra short chain PFCAs, PFBA (C4) and TFA (C2) produced nearly identical K_d_ values of 7.7 and 7.9mM respectively, but drastically different destabilization effects with ΔTm values of −29 and −3.8℃. However, both C8 PFAS congeners, PFOS and 6:2 FTSA, had nearly identical K_d_ values suggesting the presence of non-fluorinated alkyl groups point to a relatively minor role of fluorination with respect to binding affinity.

Very large Hill slopes indicative of cooperative binding was also evident for all PFAS examined. This finding is consistent with the sequential binding mechanism that has been defined for Fe^3+^ binding in the binding pocket in each of the two homologous lobular domains of hLF protein further supporting the similarity between the Fe^3+^ and PFAS binding mechanism. Our molecular docking simulations revealed that each PFAS was indeed bound within the Fe^3+^ binding sites in each lobe and that the strength of binding was generally greater at the C-lobe, a finding that further supports the idea that the mechanism of PFAS binding was similar to the sequential cooperative two-step binding mechanism of Fe^3+^ binding shown in DSF. To investigate further the similarities between Fe^3+^ and PFAS binding we evaluated the impacts of carbonate, a co-ion requisite for Fe^3+^ binding, on a subset of PFCA and PFCS congeners that act to stabilize the hLF protein structure. The stabilizing effect of carbonate was observed as a 3°C increase in the Tm of apo-LF; an effect that results from carbonate interacting with Arg463 and Thr459 of the C domain, and helix 5 and Arg121 of the N domain ^41,50^. The subsequent binding of Fe^3+^ in the C-domain is mediated by interactions with Asp395, Tyr435, Tyr528, His597, and the two oxygens of carbonate, and by two tyrosines (Tyr92, Tyr192), asparagine (Asp60), and histidine (His253) in the N-lobe ^30,51^.

Our results showed that PFAS were bound by hLF both in the presence and absence of carbonate and produced similar decreases in thermal stability. Overall binding affinity (K_d_) increased with carbon chain length and was higher for sulfonates than carboxylates (Table 1). Together, the DSF binding experiments and modeling indicate that PFAS interact with hLF through a mechanism similar to that of Fe^3+^ binding, although the effect of carbonate as a co-ion requisite was influenced by the PFAS structure; affinity generally increased with perfluorinated carbon chain length, with PFHxA, PFOS, and PFOA exhibiting higher affinity than TFA, PFBA, and PFBS (Table 1). The addition of 8mM carbonate had little effect on the maximal ΔTm for most PFAS, indicating that the structural impacts of PFAS congeners binding are similar with the notable exception of PFBA and the ultrashort-chain PFAC TFA where TFA binding affinity decreased significantly in the presence of carbonate, and PFBA had a significant impact on thermal stability of the protein with carbonate present.

The dramatic difference in both Hill slope and ΔTm for PFBA in the presence of bicarbonate suggested a non-stoichiometric binding mechanism could account for this difference. Molecular docking was performed to evaluate whether 2 molecules of PFBA could be bound in the Fe-binding pocket; indeed 2 PFBA molecules were accommodated with increased affinity through interactions with Arg465 and His597 suggesting that PFBA was substituting for carbonate as a co-ion requisite (Table 3, Table 5). This dual occupancy effect was not possible for the C4-sulfonate PFBS due to larger sulfonate headgroup which sterically limited 2 PFBS molecules in the binding pocket, a finding consistent with the minimal effects of bicarbonate on PFBS binding (Fig. 5).

**Table 5:** Single and double ligand docking of PFBA to hLF.

| Single Ligand |  |  | Double Ligand |  |  |
| --- | --- | --- | --- | --- | --- |
| # H bonds | Residues | $\Delta G$ | # H bonds | Residues | $\Delta G$ |
| 2 | Arg465, His597 | -5.4 | 3 | Thr529, Gly530, Asn640 | -7.3 |
|  |  |  | 4 | Asp395, Arg465, His597 |  |

$\Delta G$ is in kcal/mole.

**Figure 5:**
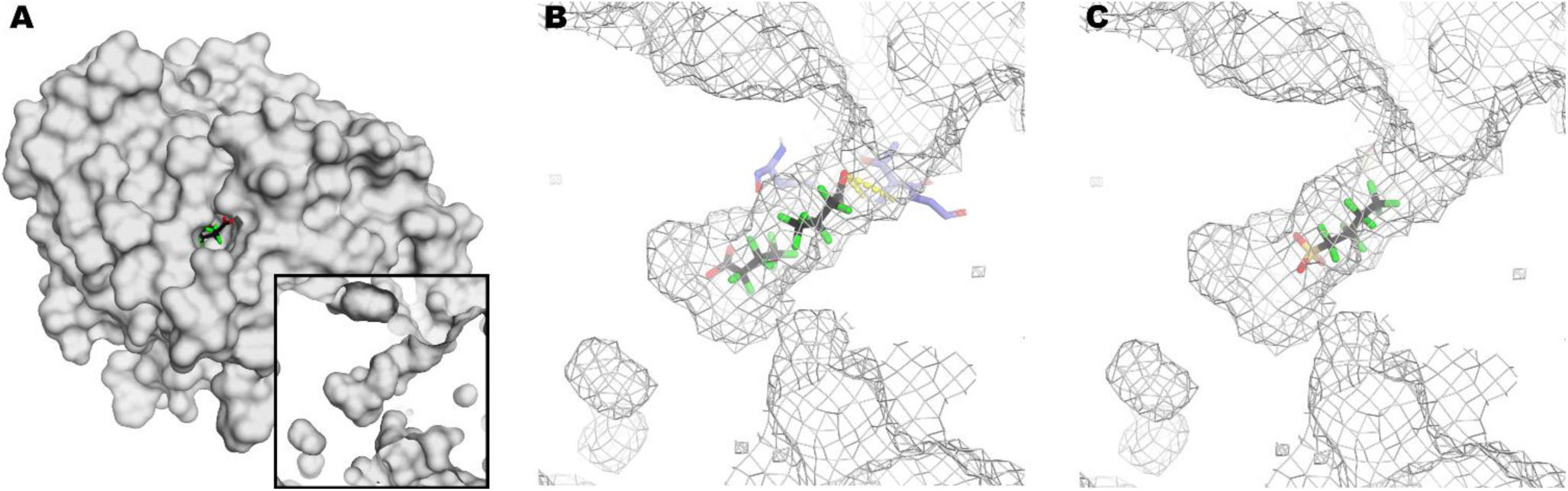
Dual Ligand Molecular Docking of PFBA and PFBS to hLF. (A) The binding pocket is shown from a down-the-pocket perspective, and after an approximately 90° rotation to reveal a cross-sectional side view of the binding cavity in the C-lobe of hLF (insert). (B) Mesh diagram of a cross-sectional side view of the binding cavity with two molecules of PFBA. Hydrogen bond interactions are depicted as dashed lines with the corresponding amino acids as stick representations and labeled. (C) Mesh diagram of the cross-sectional view of the binding cavity with PFBS.

Because bicarbonate influenced PFAS binding experimentally, additional docking was performed using holo-hLF with the Fe^3+^ ions removed while maintaining the closed C-lobe conformation. Unlike apo-hLF, blind docking revealed comparable binding to both the N- and C-lobes with similar binding energies. Most PFAS (TFA, PFBA, PFBS, PFHxA, PFHxS, PFOS, HFPO-DA, and 6:2 FTSA) occupied both bicarbonate pockets, while PFOA, PFO3DoDa, and PFDHA bound only in the N lobe. Seven PFAS (PFHxA, PFHxS, PFOA, PFOS, PFDHA, PFO3DoDa, and 6:2 FTSA) illustrated hydrogen bond interactions to His252, located in the bicarbonate binding pocket of the N-lobe. The remaining PFAS showed hydrogen bonding to either positively charged, or polar amino acids of arginine or asparagine. However, PFBS was the only PFAS to show no hydrogen bond interactions within the top 3 generated positions, indicating the binding is primarily mediated by van der Waals forces or hydrophobic effects.

To further define lobe-specific binding, molecular docking was performed separately on the C- and N-lobes of holo-hLF. PFAS binding was more favorable in the C-lobe, whereas all 11 PFAS exhibited weaker binding to the N-lobe despite minimal changes in the hydrogen bond interactions. These findings indicate that bicarbonate alters the binding location rather than completely preventing PFAS binding.

Although the presence of bicarbonate altered some hydrogen-bond interactions, PFAS continued to preferentially interact with polar or positively charged amino acids such as arginine, asparagine, or histidine. TFA was the exception, with binding driven by hydrophobic or van der Waals interactions with total loss of hydrogen bonds. These results indicate that both the apo- and holo-forms of hLF interact with PFAS through similar mechanisms, with preferential C-lobe binding followed by a conformational change that promotes N-lobe binding, consistent with the high Hill slopes observed by DSF.

Molecular docking to structurally analogous transferring proteins; human serum transferrin (h-TF), human lactoferrin (hLF), bovine lactoferrin (b-LF), *Gallus* ovotransferrin (o-TF), and human melanotransferrin (m-TF), homologous two lobed iron-binding proteins with ∼40% homology, was carried out to determine any commonality in binding mechanism across the protein family ^52^. Despite differences in physiological functions and iron-binding, molecular modeling predicted conserved PFAS-binding interactions across the transferrin family. PFAS consistently localized near the iron-binding pocket or hinge region and produced favorable binding energies. Notably, iron-coordinating residues were involved in PFAS binding, including conserved hydrogen bonding interactions with His585 of h-TF, His597 of hLF, and His592 of o-TF, suggesting PFAS may broadly disrupt transferrin function (Table 3) ^52^. These findings are consistent with our previous findings that PFAS binding is conserved across β-lactoglobulin and other calyx-domain proteins ^53^.

The DSF and molecular docking analyses provide insight into the mechanism by which PFAS interact with hLF. Across all PFAS tested, binding destabilized hLF with decreases in the thermal stability towards melting temperatures below physiological temperature, and indicate substantial disruption of hLF stability. Molecular docking demonstrated preferential binding to the C-lobe of hLF supporting the mechanism in which initial C-lobe binding promotes sequential binding to the N-lobe. Although carbonate and iron alter binding conformation, their presence does not prevent PFAS binding. This suggests that multiple conformations of hLF are susceptible to PFAS binding. Similarly, spontaneous binding was also observed across other transferrins, indicating that this binding mechanism is conserved and can contribute to PFAS-mediated disruption of immune-related proteins.

## ASSOCIATED CONTENT

### Supporting Information

The following files are available free of charge.

brief description (file type, i.e., PDF)

brief description (file type, i.e., PDF)

## AUTHOR INFORMATION

### Author Contributions

The manuscript was written through contributions of all authors. All authors have given approval to the final version of the manuscript. ‡These authors contributed equally.

### Funding Sources

The research reported herein was supported by the National Institute of Environmental Health Sciences of the National Institutes of Health under award numbers P01ES035542, P42ES031009, P30ES025128, T3ES007046, National Science Foundation award number OCE-2414792, and a fellowship awarded to ZM from NSF award OISE-2420222. The content is solely the responsibility of the authors and does not necessarily represent the official views of the National Institutes of Health nor the National Science Foundation.

## ABBREVIATIONS

hLF: Lactoferrin
DF: Deferiprone
DSF: Differential Scanning Fluorimetry
DSC: Differential Scanning Calorimetry
o-TF: Ovotransferrin
m-TF: Melanotransferrin
b-LF: Bovine Lactoferrin
h-TF: Human Serum Transferrin

## Notes

### Competing Interest Statement

The authors have declared no competing interest.

## REFERENCES

(1) Routti, H.; Krafft, B. A.; Herzke, D.; Eisert, R.; Oftedal, O. Perfluoroalkyl Substances Detected in the World’s Southernmost Marine Mammal, the Weddell Seal ( Leptonychotes Weddellii). Environ. Pollut. 2015, 197, 62–67. 10.1016/j.envpol.2014.11.026.

(2) Olsen, G. W.; Mair, D. C.; Lange, C. C.; Harrington, L. M.; Church, T. R.; Goldberg, C. L.; Herron, R. M.; Hanna, H.; Nobiletti, J. B.; Rios, J. A.; Reagen, W. K.; Ley, C. A. Per- and Polyfluoroalkyl Substances (PFAS) in American Red Cross Adult Blood Donors, 2000– 2015. Environ. Res. 2017, 157, 87–95. 10.1016/j.envres.2017.05.013.

(3) Koch, A.; Jonsson, M.; Yeung, L. W. Y.; Kärrman, A.; Ahrens, L.; Ekblad, A.; Wang, T. Quantification of Biodriven Transfer of Per- and Polyfluoroalkyl Substances from the Aquatic to the Terrestrial Environment via Emergent Insects. Environ. Sci. Technol. 2021, 55 (12), 7900–7909. 10.1021/acs.est.0c07129.

(4) Birgersson, L.; Jouve, J.; Jönsson, E.; Asker, N.; Andreasson, F.; Golovko, O.; Ahrens, L.; Sturve, J. Thyroid Function and Immune Status in Perch (Perca Fluviatilis) from Lakes Contaminated with PFASs or PCBs. Ecotoxicol. Environ. Saf. 2021, 222, 112495. 10.1016/j.ecoenv.2021.112495.

(5) Fair, P. A.; Romano, T.; Schaefer, A. M.; Reif, J. S.; Bossart, G. D.; Houde, M.; Muir, D.; Adams, J.; Rice, C.; Hulsey, T. C.; Peden-Adams, M. Associations between Perfluoroalkyl Compounds and Immune and Clinical Chemistry Parameters in Highly Exposed Bottlenose Dolphins (Tursiops Truncatus). Environ. Toxicol. Chem. 2013, 32 (4), 736–746. 10.1002/etc.2122.

(6) Custer, C. M.; Custer, T. W.; Schoenfuss, H. L.; Poganski, B. H.; Solem, L. Exposure and Effects of Perfluoroalkyl Compounds on Tree Swallows Nesting at Lake Johanna in East Central Minnesota, USA. Reprod. Toxicol. 2012, 33 (4), 556–562. 10.1016/j.reprotox.2011.01.005.

(7) Custer, C. M.; Custer, T. W.; Dummer, P. M.; Etterson, M. A.; Thogmartin, W. E.; Wu, Q.; Kannan, K.; Trowbridge, A.; McKann, P. C. Exposure and Effects of Perfluoroalkyl Substances in Tree Swallows Nesting in Minnesota and Wisconsin, USA. Arch. Environ. Contam. Toxicol. 2014, 66 (1), 120–138. 10.1007/s00244-013-9934-0.

(8) Domingo, J. L. A Review of the Occurrence and Distribution of Per- and Polyfluoroalkyl Substances (PFAS) in Human Organs and Fetal Tissues. Environ. Res. 2025, 272, 121181. 10.1016/j.envres.2025.121181.

(9) Liu, D.; Yan, S.; Wang, P.; Chen, Q.; Liu, Y.; Cui, J.; Liang, Y.; Ren, S.; Gao, Y. Perfluorooctanoic Acid (PFOA) Exposure in Relation to the Kidneys: A Review of Current Available Literature. Front. Physiol. 2023, 14. 10.3389/fphys.2023.1103141.

(10) Fenton, S. E.; Ducatman, A.; Boobis, A.; DeWitt, J. C.; Lau, C.; Ng, C.; Smith, J. S.; Roberts, S. M. Per- and Polyfluoroalkyl Substance Toxicity and Human Health Review: Current State of Knowledge and Strategies for Informing Future Research. Environ. Toxicol. Chem. 2021, 40 (3), 606–630. 10.1002/etc.4890.

(11) Lu, Y.; Guan, R.; Zhu, N.; Hao, J.; Peng, H.; He, A.; Zhao, C.; Wang, Y.; Jiang, G. A Critical Review on the Bioaccumulation, Transportation, and Elimination of per- and Polyfluoroalkyl Substances in Human Beings. Crit. Rev. Environ. Sci. Technol. 2024, 54 (2), 95–116. 10.1080/10643389.2023.2223118.

(12) Moro, G.; Liberi, S.; Vascon, F.; Linciano, S.; De Felice, S.; Fasolato, S.; Foresta, C.; De Toni, L.; Di Nisio, A.; Cendron, L.; Angelini, A. Investigation of the Interaction between Human Serum Albumin and Branched Short-Chain Perfluoroalkyl Compounds. Chem. Res. Toxicol. 2022, 35 (11), 2049–2058. 10.1021/acs.chemrestox.2c00211.

(13) Forsthuber, M.; Kaiser, A. M.; Granitzer, S.; Hassl, I.; Hengstschläger, M.; Stangl, H.; Gundacker, C. Albumin Is the Major Carrier Protein for PFOS, PFOA, PFHxS, PFNA and PFDA in Human Plasma. Environ. Int. 2020, 137, 105324. 10.1016/j.envint.2019.105324.

(14) Birchfield, A. S.; Musayev, F. N.; Castillo, A. J.; Zorn, G.; Fuglestad, B. Broad PFAS Binding with Fatty Acid Binding Protein 4 Is Enabled by Variable Binding Modes. BioRxiv Prepr. Serv. Biol. 2025, 2025.01.10.632451. 10.1101/2025.01.10.632451.

(15) Mastromarino, P.; Capobianco, D.; Campagna, G.; Laforgia, N.; Drimaco, P.; Dileone, A.; Baldassarre, M. E. Correlation between Lactoferrin and Beneficial Microbiota in Breast Milk and Infant’s Feces. BioMetals 2014, 27 (5), 1077–1086. 10.1007/s10534-014-9762-3.

(16) Rai, D.; Adelman, A. S.; Zhuang, W.; Rai, G. P.; Boettcher, J.; Lönnerdal, B. Longitudinal Changes in Lactoferrin Concentrations in Human Milk: A Global Systematic Review,. Crit. Rev. Food Sci. Nutr. 2014, 54 (12), 1539–1547. 10.1080/10408398.2011.642422.

(17) Cacho, N. T.; Lawrence, R. M. Innate Immunity and Breast Milk. Front. Immunol. 2017, 8, 584. 10.3389/fimmu.2017.00584.

(18) Moles, L.; Manzano, S.; Fernández, L.; Montilla, A.; Corzo, N.; Ares, S.; Rodríguez, J. M.; Espinosa-Martos, I. Bacteriological, Biochemical, and Immunological Properties of Colostrum and Mature Milk From Mothers of Extremely Preterm Infants. J. Pediatr. Gastroenterol. Nutr. 2015, 60 (1), 120–126. 10.1097/MPG.0000000000000560.

(19) Trend, S.; Strunk, T.; Hibbert, J.; Kok, C. H.; Zhang, G.; Doherty, D. A.; Richmond, P.; Burgner, D.; Simmer, K.; Davidson, D. J.; Currie, A. J. Antimicrobial Protein and Peptide Concentrations and Activity in Human Breast Milk Consumed by Preterm Infants at Risk of Late-Onset Neonatal Sepsis. PLoS ONE 2015, 10 (2), e0117038. 10.1371/journal.pone.0117038.

(20) Stănciuc, N.; Aprodu, I.; Râpeanu, G.; Van Der Plancken, I.; Bahrim, G.; Hendrickx, M. Analysis of the Thermally Induced Structural Changes of Bovine Lactoferrin. J. Agric. Food Chem. 2013, 61 (9), 2234–2243. 10.1021/jf305178s.

(21) Andersen, B. F.; Baker, H. M.; Morris, G. E.; Rumball, S. V.; Baker, E. N. Apolactoferrin Structure Demonstrates Ligand-Induced Conformational Change in Transferrins. Nature 1990, 344 (6268), 784–787. 10.1038/344784a0.

(22) Haridas, M.; Anderson, B. F.; Baker, E. N. Structure of Human Diferric Lactoferrin Refined at 2.2 Å Resolution. Acta Crystallogr. D Biol. Crystallogr. 1995, 51 (5), 629–646. 10.1107/S0907444994013521.

(23) Jameson, G. B.; Anderson, B. F.; Norris, G. E.; Thomas, D. H.; Baker, E. N. Structure of Human Apolactoferrin at 2.0 Å Resolution. Refinement and Analysis of Ligand-Induced Conformational Change. Acta Crystallogr. D Biol. Crystallogr. 1998, 54 (6), 1319–1335. 10.1107/S0907444998004417.

(24) Farnaud, S.; Evans, R. W. Lactoferrin—a Multifunctional Protein with Antimicrobial Properties. Mol. Immunol. 2003, 40 (7), 395–405. 10.1016/S0161-5890(03)00152-4.

(25) Aisen, P.; Leibman, A. Lactoferrin and Transferrin: A Comparative Study. Biochim. Biophys. Acta BBA - Protein Struct. 1972, 257 (2), 314–323. 10.1016/0005-2795(72)90283-8.

(26) Baker, E. N. Structure and Reactivity of Transferrins. In Advances in Inorganic Chemistry; Sykes, A. G., Ed.; Academic Press, 1994; Vol. 41, pp 389–463. 10.1016/S0898-8838(08)60176-2.

(27) Brisson, G.; Britten, M.; Pouliot, Y. Effect of Iron Saturation on the Recovery of Lactoferrin in Rennet Whey Coming from Heat-Treated Skim Milk. J. Dairy Sci. 2007, 90 (6), 2655– 2664. 10.3168/jds.2006-725.

(28) Baker, E. N.; Baker, H. M.; Kidd, R. D. Lactoferrin and Transferrin: Functional Variations on a Common Structural Framework. Biochem. Cell Biol. 2002, 80 (1), 27–34. 10.1139/o01-153.

(29) Gerstein, M.; Anderson, B. F.; Norris, G. E.; Baker, E. N.; Lesk, A. M.; Chothia, C. Domain Closure in Lactoferrin: Two Hinges Produce a See-Saw Motion Between Alternative Close-Packed Interfaces. J. Mol. Biol. 1993, 234 (2), 357–372. 10.1006/jmbi.1993.1592.

(30) Dyrda-Terniuk, T.; Pomastowski, P. The Multifaceted Roles of Bovine Lactoferrin: Molecular Structure, Isolation Methods, Analytical Characteristics, and Biological Properties. J. Agric. Food Chem. 2023, 71 (51), 20500–20531. 10.1021/acs.jafc.3c06887.

(31) Baker, E. N.; Baker, H. M. A Structural Framework for Understanding the Multifunctional Character of Lactoferrin. Biochimie 2009, 91 (1), 3–10. 10.1016/j.biochi.2008.05.006.

(32) Starnes, H. M.; Jackson, T. W.; Rock, K. D.; Belcher, S. M. Quantitative Cross-Species Comparison of Serum Albumin Binding of per- and Polyfluoroalkyl Substances from Five Structural Classes. Toxicol. Sci. 2024, 199 (1), 132–149. 10.1093/toxsci/kfae028.

(33) Starnes, H. M.; Belcher, S. M. Protocol for Evaluating Protein-Polyfluoroalkyl Substances *in Vitro* Using Differential Scanning Fluorimetry. STAR Protoc. 2024, 5 (4), 103386. 10.1016/j.xpro.2024.103386.

(34) Jackson, T. W.; Scheibly, C. M.; Polera, M. E.; Belcher, S. M. Rapid Characterization of Human Serum Albumin Binding for Per- and Polyfluoroalkyl Substances Using Differential Scanning Fluorimetry. Environ. Sci. Technol. 2021, 55 (18), 12291–12301. 10.1021/acs.est.1c01200.

(35) Eberhardt, J.; Santos-Martins, D.; Tillack, A. F.; Forli, S. AutoDock Vina 1.2.0: New Docking Methods, Expanded Force Field, and Python Bindings. J. Chem. Inf. Model. 2021, 61 (8), 3891–3898. 10.1021/acs.jcim.1c00203.

(36) Trott, O.; Olson, A. J. AutoDock Vina: Improving the Speed and Accuracy of Docking with a New Scoring Function, Efficient Optimization and Multithreading. J. Comput. Chem. 2010, 31 (2), 455–461. 10.1002/jcc.21334.

(37) Schrodinger L.L.C. PyMOL, 2017. https://pymol.org.

(38) Morris, G. M.; Huey, R.; Lindstrom, W.; Sanner, M. F.; Belew, R. K.; Goodsell, D. S.; Olson, A. J. AutoDock4 and AutoDockTools4: Automated Docking with Selective Receptor Flexibility. J. Comput. Chem. 2009, 30 (16), 2785–2791. 10.1002/jcc.21256.

(39) Mizutani, K.; Muralidhara, B. K.; Yamashita, H.; Tabata, S.; Mikami, B.; Hirose, M. Anion-Mediated Fe3+ Release Mechanism in Ovotransferrin C-Lobe. J. Biol. Chem. 2001, 276 (38), 35940–35946. 10.1074/jbc.M102590200.

(40) Hayashi, K.; Longenecker, K. L.; Liu, Y.-L.; Faust, B.; Prashar, A.; Hampl, J.; Stoll, V.; Vivona, S. Complex of Human Melanotransferrin and SC57.32 Fab Fragment Reveals Novel Interdomain Arrangement with Ferric N-Lobe and Open C-Lobe. Sci. Rep. 2021, 11 (1), 566. 10.1038/s41598-020-79090-8.

(41) Sharma, S.; Jasti, J.; Kumar, J.; Mohanty, A. K.; Singh, T. P. Crystal Structure of a Proteolytically Generated Functional Monoferric C-Lobe of Bovine Lactoferrin at 1.9 Å Resolution. J. Mol. Biol. 2003, 331 (2), 485–496. 10.1016/S0022-2836(03)00717-4.

(42) Yang, N.; Zhang, H.; Wang, M.; Hao, Q.; Sun, H. Iron and Bismuth Bound Human Serum Transferrin Reveals a Partially-Opened Conformation in the N-Lobe. Sci. Rep. 2012, 2 (1), 999. 10.1038/srep00999.

(43) Vivoli, M.; Novak, H. R.; Littlechild, J. A.; Harmer, N. J. Determination of Protein-Ligand Interactions Using Differential Scanning Fluorimetry. J. Vis. Exp. JoVE 2014, No. 91, 51809. 10.3791/51809.

(44) Savitzky, Abraham.; Golay, M. J. E. Smoothing and Differentiation of Data by Simplified Least Squares Procedures. Anal. Chem. 1964, 36 (8), 1627–1639. 10.1021/ac60214a047.

(45) von Ehrenstein, O. S.; Fenton, S. E.; Kato, K.; Kuklenyik, Z.; Calafat, A. M.; Hines, E. P. Polyfluoroalkyl Chemicals in the Serum and Milk of Breastfeeding Women. Reprod. Toxicol. 2009, 27 (3), 239–245. 10.1016/j.reprotox.2009.03.001.

(46) Blake, B. E.; Fenton, S. E. Early Life Exposure to Per- and Polyfluoroalkyl Substances (PFAS) and Latent Health Outcomes: A Review Including the Placenta as a Target Tissue and Possible Driver of Peri- and Postnatal Effects. Toxicology 2020, 443, 152565. 10.1016/j.tox.2020.152565.

(47) LaKind, J. S.; Naiman, J.; Verner, M.-A.; Lévêque, L.; Fenton, S. Per- and Polyfluoroalkyl Substances (PFAS) in Breast Milk and Infant Formula: A Global Issue. Environ. Res. 2023, 219, 115042. 10.1016/j.envres.2022.115042.

(48) Grandjean, P.; Andersen, E. W.; Budtz-Jørgensen, E.; Nielsen, F.; Mølbak, K.; Weihe, P.; Heilmann, C. Serum Vaccine Antibody Concentrations in Children Exposed to Perfluorinated Compounds. JAMA 2012, 307 (4), 391–397. 10.1001/jama.2011.2034.

(49) Herich, R.; Levkut, M.; Bomba, A.; Gancarcíková, S.; Nemcová, R. Differences in the Development of the Small Intestine between Gnotobiotic and Conventionally Bred Piglets. Berl. Munch. Tierarztl. Wochenschr. 2004, 117 (1–2), 46–51.

(50) Baker, E. N.; Baker, H. M. Lactoferrin. Cell. Mol. Life Sci. 2005, 62 (22), 2531. 10.1007/s00018-005-5368-9.

(51) Paramasivan, M.; Karthikeyan, S.; Sharma, S.; Sharma, A. K.; Paramasivam, M.; Yadav, S.; Srinivasan, A.; Singh, T. P. Structural Variability and Functional Convergence in Lactoferrins. Curr. Sci. 1999, 77 (2), 241–255.

(52) Halbrooks, P. J.; Giannetti, A. M.; Klein, J. S.; Björkman, P. J.; Larouche, J. R.; Smith, V. C.; MacGillivray, R. T. A.; Everse, S. J.; Mason, A. B. Composition of pH-Sensitive Triad in C-Lobe of Human Serum Transferrin. Comparison to Sequences of Ovotransferrin and Lactoferrin Provides Insight into Functional Differences in Iron Release. Biochemistry 2005, 44 (47), 15451–15460. 10.1021/bi0518693.

(53) McLean, Z. S.; Thomas, M. E.; Belcher, S. M. β-Lactoglobulin – PFAS Binding Interactions Identifies the Calyx Domain as a Determinant of Contaminated Milk Exposure and the Calycin Protein Family as Potential Mediators of PFAS Toxicity. Toxicol. Sci. 2025, kfaf178. 10.1093/toxsci/kfaf178.

