## Supplementary Material for "Human Lactoferrin is a Novel PFAS Target: Implications for Neo-natal Immune Function and Protein Stability"

S1: Grid Box Dimensions and Coordinates for Molecular Docking.

| Protein | PDB ID | Species | X (Å) | Y (Å) | Z (Å) | X center | Y center | Z center |
|---|---|---|---|---|---|---|---|---|
| Apo-hLF | 1CB6 | Human | 86 | 80 | 64 | 18.901 | 14.610 | -24.045 |
| Apo-hLF C lobe | | | 46 | 46 | 78 | 40.812 | 30.869 | -16.036 |
| Apo-hLF N lobe | | | 60 | 56 | 66 | 3.981 | 4.225 | -27.853 |
| Holo-hLF | 1LFG | Human | 86 | 80 | 64 | -5.038 | 23.058 | 10.537 |
| Holo-hLF C lobe | | | 46 | 58 | 56 | 15.374 | 29.831 | 20.749 |
| Holo-hLF N lobe | | | 60 | 56 | 66 | -20.361 | -1.361 | 36.111 |
| Ovotransferrin | 1IQ7 | Gallus | 58 | 52 | 62 | 64.587 | -17.909 | 53.790 |
| Melanotransferrin | 6XR0 | Human | 50 | 66 | 60 | 7.745 | -6.596 | 1.285 |
| Serum Transferrin | 4H0W | Human | 50 | 66 | 60 | 7.745 | -6.596 | 1.285 |
| Bovine Lactoferrin | 1NKX | Bovine | 60 | 54 | 56 | 3.950 | 0.962 | 13.951 |

S2: (A) Normalized derivative fluorescence spectrum of LF with increasing additions of EDTA. The Tm values for LF are denoted by dashed lines. (B) Concentration response curve of LF with increasing additions of EDTA. Error bars indicate the standard error of the mean.

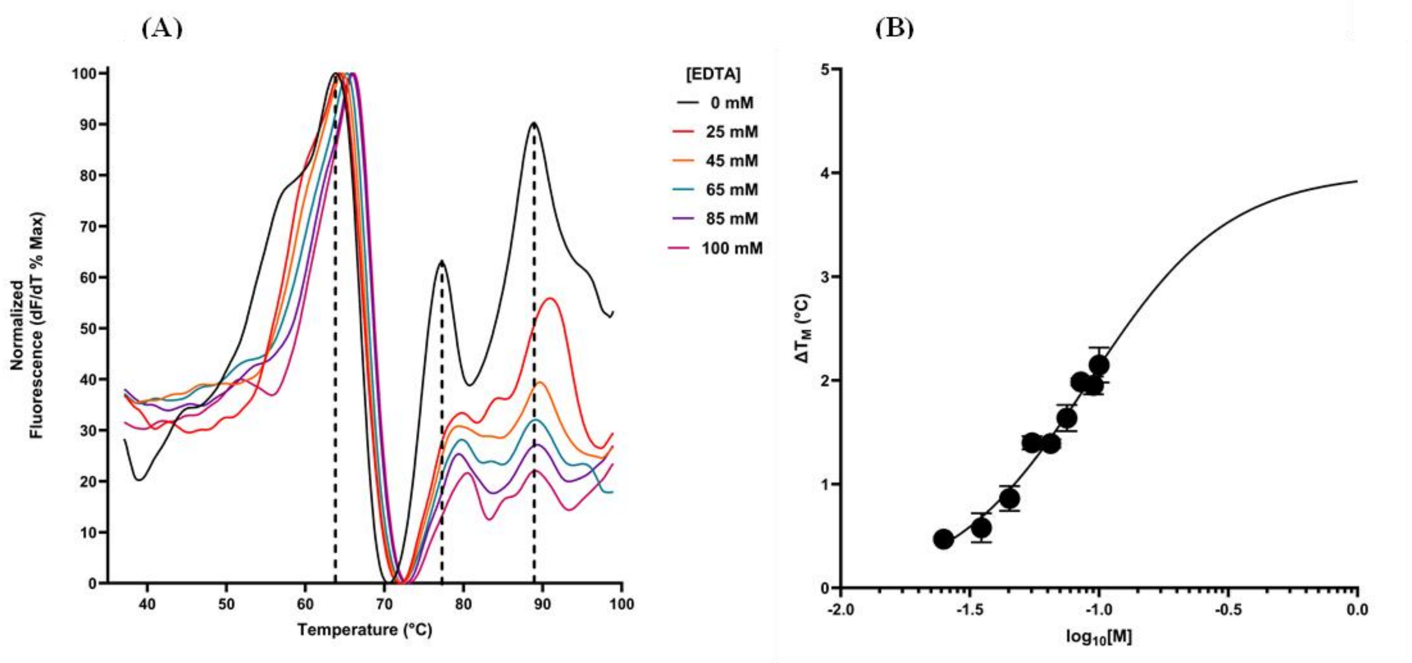


S3: (A) Normalized derivative fluorescence of LF with increasing MeOH (v/v%). The Tm values for LF are indicated by the dotted lines. (B) Concentration response curve of LF with increasing additions of MeOH.  Error bars indicate the standard error of the mean.

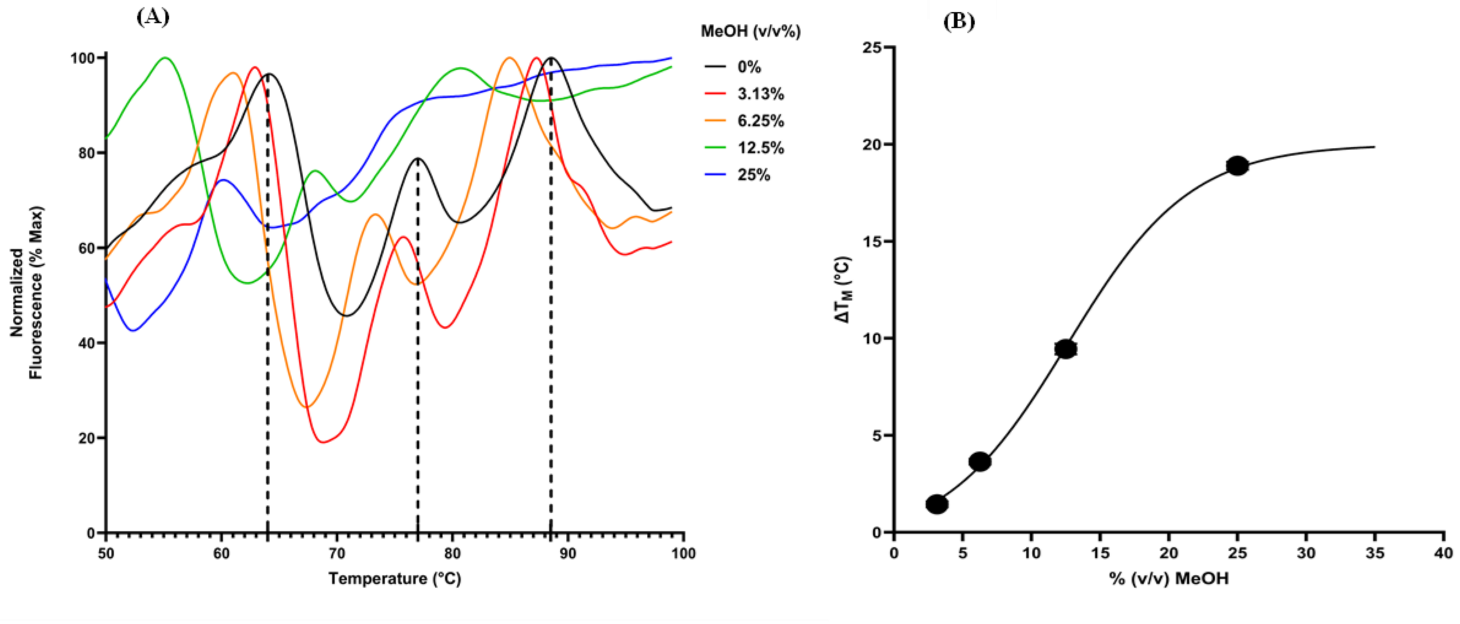

S4: Crystal structure similarity between transferrin-family proteins and apo-hLF.

| **Protein** | **PDB ID** | **RMSD** |
|---|---|---|
| Ovotransferrin | 1IQ7 | 0.778 |
| Melanotransferrin | 6XR0 | 1.513 |
| Bovine Lactoferrin | 1NKX | 4.520 |
| Serum Transferrin | 4H0W | 7.167 |

*RMSD: root mean square deviation. The RMSD is in Å, and was derived from the differences between apo-hLF (1CB6) and each of the transferrins.*
